# Cell-specific regulation of TCA cycle and cell division related genes at roots following aluminium exposure

**DOI:** 10.1101/2025.10.07.680940

**Authors:** Welder Alves Silva, Auxiliadora Oliveira Martins, Thiago Wakin, Marcelle Ferreira Silva, Joao Antonio Siqueira, David B. Medeiros, Dimas M. Ribeiro, Alisdair R. Fernie, Adriano Nunes-Nesi, Wagner L. Araújo

**Author notes:** Correspondence Wagner L. Araújo Departamento de Biologia Vegetal Universidade Federal de Viçosa 36570-900 Viçosa, Minas Gerais, Brazi.

## Abstract

Aluminium (Al) toxicity is a critical factor that limits plant growth in acidic soils (pH < 5.5) worldwide. Intracellular Al can bind to various biomolecules and significantly affect gene expression, protein biosynthesis, and cell membrane integrity, ultimately inhibiting nutrients and water uptake. This leads to the arrest of root cell divisions and growth. Organic acids (OAs) produced in mitochondrial-related reactions are strongly linked to Al tolerance, as they bind to Al and neutralize its toxic effects. Recently, the manipulation of root cell divisions through DNA checkpoints has been proposed as an alternative to increase Al tolerance. In this context, molecular interactions between these two mechanisms could potentially enhance plant tolerance to Al toxicity. Here, we investigate this topic using a translatome approach, which enables specific analyses of root cell types (namely, whole root, the quiescent centre region, and the cortical region). This approach offers a promising tool for unravelling how mitochondrial metabolism, particularly the TCA cycle, and the DNA damage response (DDR) pathway are interconnected in Al-mitigation process. Our results revealed that the TCA cycle is induced in differentiated cells, with no significant alteration in meristematic cells, while genes associated with cell cycle progression and DNA checkpoints show differential expression across the distinct cell types. Collectively, our data contribute to understand cell-specific internal detoxification mechanism related to Al tolerance in plants. This knowledge could assist crop breeding by providing a better understanding of the cellular physiological responses to Al toxicity.

## INTRODUCTION

Acidic soil conditions (pH < 5.5) are distributed worldwide, and most of them display chemical properties leading to the release of soluble aluminium (Al) cations. Intracellular Al can bind to various biomolecules, significantly affecting gene expression, protein biosynthesis, and the integrity of cell membranes (Kochian et al., 2004; 2015). The disruption of these processes contributes to reducing the uptake of water and nutrients, which eventually arrests root cell divisions and growth (Nunes-Nesi et al., 2014; Siqueira et al., 2022). The most discussed mechanism that plants use to mitigate Al toxicity involves its neutralization in the intra or extracellular space (Nunes-Nesi et al., 2014; Kochian et al., 2015). Additionally, organic acids (OAs) produced in mitochondrial-related reactions can be exuded from root cells into the rhizosphere, where these compounds bind to Al and neutralize this metal (Nunes-Nesi et al., 2014; Kochian et al., 2015). Several studies have manipulated genes involved in OA exudation to minimize the negative impacts of Al toxicity on crop yield, which indicated certain limitations to engineering crops solely through this mechanism (Ofoe et al., 2022b). Accordingly, even Al-sensitive genotypes release high amounts of malate and citrate into the soil, whereas Al tolerance is seemingly intimately supported by endogenous OAs that contribute to intracellular detoxification (Ma et al., 1997, Ma et al., 1998, Ishikawa et al., 2000, Piñeros et al., 2005, Zheng et al., 2005). Since OAs are mainly produced through the mitochondrial tricarboxylic acid (TCA) cycle, and its intermediaries have distinct impacts on root phenotypes, the significance of these compounds for specific genotypes should be taken into consideration (Zhang et al., 2023). It has been revealed that the genetic background of genotypes is crucial in orchestrating OA exudation mechanisms (Melo et al., 2019), indicating additional effects for the manipulation of OA exudation.

Over the last two decades, the potential for manipulating root cell divisions through DNA checkpoints to increase Al tolerance has been suggested (Rounds and Larsen, 2008; Nezames et al., 2012; Sjogren et al., 2015; Chen et al., 2019). Briefly, due to the presence of phosphate in nucleotides, DNA is negatively charged and a major target for binding by Al cations (Hu et al., 2016; Siqueira et al., 2022). Upon the interaction of Al with DNA, single and double-strand breaks can be generated, resulting in DNA damage that arrests cell divisions (Hu et al., 2016). When DNA damage occurs, early responses involve recognizing the damaged DNA sites, where sensor protein complexes detect the damage and, depending on the type of damage, can activate the protein kinases ATAXIA TELANGIECTASIA MUTATED (ATM) and ATM- AND RAD3-RELATED (ATR) (Maréchal, A. and Zou, L., 2013). Both ATM and ATR kinases activate the transcription factor SUPPRESSOR OF GAMMA-RESPONSE 1 (SOG1) upon DNA damage (Yoshiyama et al., 2009; Yoshiyama et al., 2013; Sjogren et al., 2015). After activation, SOG1 mediates the induction of several genes involved in DNA repair, cell cycle arrest, and ultimately programmed cell death (Fulcher and Sablowski, 2009; Furukawa et al., 2010; Adachi et al., 2011). The cell cycle is interrupted for DNA repair to prevent the transmission of mutations to daughter cells. SOG1 induces the genes *SIAMESE/SIAMESE-RELATED 5* (*SMR5*) and *SMR7*, which in turn inhibit *CYCLIN-DEPENDENT KINASE* (*CDK*), blocking cell cycle progression (Yi et al., 2014). Therefore, the loss-of-function mutants *atm*, *atr*, and *sog1* were characterized to maintain cell cycle progression under Al toxicity, supporting root elongation during this stressful condition (Chen et al., 2017). Similarly to the OA mechanism, these genetic manipulations were insufficient to mediate plant survival under high Al concentrations (Chen et al., 2019). In this context, molecular interactions between these mechanisms are likely capable of improving plant tolerance to Al toxicity.

Plants have developed complex mechanisms to regulate gene expression in specific root cell types to cope with dynamic environments. Root meristems display a higher degree of gene expression conservation across different species than other cell types, whereas cell fate regulators present in root hair cells appear to control abiotic stress responses (Maher et al., 2018; Kajala et al., 2021). Although genes encoding organic anion transporters have been established as targets for increasing Al tolerance, other recently discovered mechanisms provide additional options for manipulating mitochondrial metabolism and cell cycle progression, increasing Al tolerance (for further details, see Siqueira et al., 2022). Here, by using a translatome, a profile of the translational status of gene expression within cells, our results revealed how the DNA damage response pathway and mitochondrial metabolism are differentially reprogrammed in specific root cell types. We demonstrated that genes encoding components of the TCA cycle are induced in differentiated cells, with no alteration in meristematic cells. Conversely, genes associated with cell cycle progression and DNA checkpoints show differential expression across the distinct cell types investigated here. Altogether, our results provide a high-resolution model that aids genetic improvement to increase crop tolerance to Al, especially in acidic soils.

## MATERIALS AND METHODS

### Plant material and growth conditions

*Arabidopsis thaliana* seedlings (Columbia background, Col-0) expressing the *p:GFP-FLAG-RPL18* construct (*p*: promoter; *GFP*: green fluorescent protein; *FLAG-RPL18*: FLAG-epitope tagged ribosomal protein L18) (Mustroph et al., 2009) were used. Individual promoters were used to direct the expression of the transgene, namely: Cauliflower mosaic virus 35S (*p35S:GFP-FLAG-RPL18*), almost constitutive; plastid endopeptidase (*pPEP:GFP-FLAG-RPL18*), specific to the root cortex of the elongation and maturation zones; SCARECROW (*pSCR:GFP-FLAG-RPL18*), specific to the quiescent centre and root endodermis (Supplementary Figure 1).

Seeds were sterilised in 70% ethanol and 2.5% (v/v) sodium hypochlorite and then distributed in Petri dishes containing growth medium [0.43% (*w/v*) MS medium and 0.7% (*w/v*) agar, pH 4.0 (Murashige and Skoog, 1962)] and placed in the dark at a low temperature (4°C) for 48 hours. After this period, the seedlings were kept for ten days in a controlled environment with a photoperiod of 16/8 hours (light/dark), a temperature of 22/20°C (light/dark), and light intensity of 150 μmols of photons m^-2^s^-1^. During this period, the seedlings were subjected to two treatments: the absence (0 μM) and the presence (50 μM) of Al (AlCl_3_ - Aluminium chloride hexahydrate, 99% - Honeywell^®^) in the growth medium. After ten days, whole roots from the *p35S:GFP-FLAG-RPL18* line and whole seedlings from the *pPEP:GFP-FLAG-RPL18* and *pSCR:GFP-FLAG-RPL18* lines were collected at the start of the light period and promptly stored at −80°C until metabolic profiling and translatome analyses were carried out.

### Confirmation of transgene expression

Confirmation of the *p:GFP-FLAG-RPL18* constructs expression was carried out using a Zeiss LSM510 META Laser Scanning Confocal Microscope. To do this, root apices excised from 7-day-old *Arabidopsis thaliana* seedlings grown vertically on solid growth medium [0.43% (*w/v*) MS medium and 0.7% (*w/v*) agar, pH 4.0 (Murashige and Skoog, 1962)] were used. The wavelengths used for excitation and emission of the GFP were 488 and 510 nm, respectively. Representative images were obtained using ZEISS ZEN 3.1 software (Supplementary Figure 2).

### Root growth

Seeds were sterilised in 70% ethanol and 2.5% (v/v) sodium hypochlorite and then distributed in square Petri dishes (12 x 12 cm) containing solid growth medium [0.43% (*w/v*) MS medium and 0.7% (*w/v*) agar, pH 4.0 (Murashige and Skoog, 1962)] and placed in the dark at 4°C for 48 hours. After stratification, the seedlings were kept for ten days in a controlled environment with a photoperiod of 16/8 hours (light/dark), a temperature of 22/20°C (light/dark) and light intensity of 150 μmols of photons m^-2^s^-1^. Seedlings were subjected to two treatments: absence (0 μM) and presence (50 μM) of Al (AlCl_3_) in the growth medium. To assess root growth, images were registered using the HP Scanjet G2410 scanner and processed using the ImageJ software.

### Metabolite profile

The extraction of metabolites was carried out following a protocol based on gas chromatography coupled with mass spectrometry (GC-MS) (Lisec et al., 2006). For this, approximately 25 mg of homogenised root tissue was extracted in 1,400 μL of water, methanol and chloroform (1:2.5:1, v/v/v). Next, 60 μL of internal standard (0.2 mg ribitol mL^-1^ water) was added as a quantification standard. For derivatisation, 200 μL of the top layer (polar phase) was dried under vacuum. Thus, 40 μL of methoxyamine was added to each sample, followed by the addition of 70 μL of the N-methyl-N-(trimethylsilyl)trifluoroacetamide and fatty acid methyl ester standards. The chromatograms and mass spectra were evaluated using TAGFINDER software (Luedemann et al., 2008). The metabolites were identified by comparison with databases of authentic standards (Kopka et al., 2005; Schauer et al., 2005). The identification and annotation of detected peaks followed previously described recommendations (Fernie et al., 2011).

### Gene expression

To analyse the cell-specific translatome, the translating ribosome affinity purification (TRAP) method (Mustroph et al., 2013) was used. Initially, using a mix of reagents called polysome extraction buffer, the polysomes were extracted from the homogenised root tissue. Then, using previously washed ANTI-FLAG agarose beads, another mix of reagents called washing buffer and the 3x FLAG elution peptide, the previously extracted polysomes were immunopurified. The RNA contained in the immunopurified polysomes was extracted using the RNeasy Plant Mini Kit (QIAGEN) and amplified using the Complete Whole Transcriptome Amplification Kit performed according to the manufacturer’s recommendations. In addition to RNA amplification, the Complete Whole Transcriptome Amplification Kit also synthesised the cDNA. The qRT-PCR reactions were carried out on the QuantStudio™ 1 Real-Time PCR System, Applied Biosystems, CA, USA. Using the synthesised cDNA, gene-specific primers (Supplementary Table 1) and the GoTaq® qPCR Master Mix (Promega Corporation, WI, USA) according to the following manufacturer’s recommendations. The gene encoding the F-box protein in *Arabidopsis thaliana* (At5g15710) was used as an endogenous control to normalise the qRT-PCR values. All primers used are specified in Supplementary Table 1.

### Statistical analyses

Statistical analyses were carried out using the algorithms in the Microsoft Excel program. The data relating to the analyses of root growth and gene expression were expressed as the mean of five and three repetitions, respectively, ± standard error. The data relating to the metabolite profile, expressed as the mean of four replicates, was normalised in relation to the values found for each genotype in the absence of aluminium and log_2_-transformed for representation on the heat map. The graphs were produced using SigmaPlot software. An asterisk (*) indicates a statistically significant difference between treatments within each genotype (Student *t*-test, \**p*<0.05).

## RESULTS

### Aluminium inhibited root elongation in *Arabidopsis thaliana*

As expected, a significant decrease in root growth was observed in *Arabidopsis thaliana* wild type (Columbia, Col-0) following exposure to 50 μM of aluminium (Al) after 10 days (Figure 1), which is a well-known symptom of Al toxicity (Kochian et al., 2015). While most studies on Al tolerance have focused on exudation mechanisms, there are certain limitations to engineering crops solely based on this mechanism. Therefore, our study aimed to investigate root internal detoxification mechanisms at a cellular specific level using a translatome approach.

**Figure 1.**
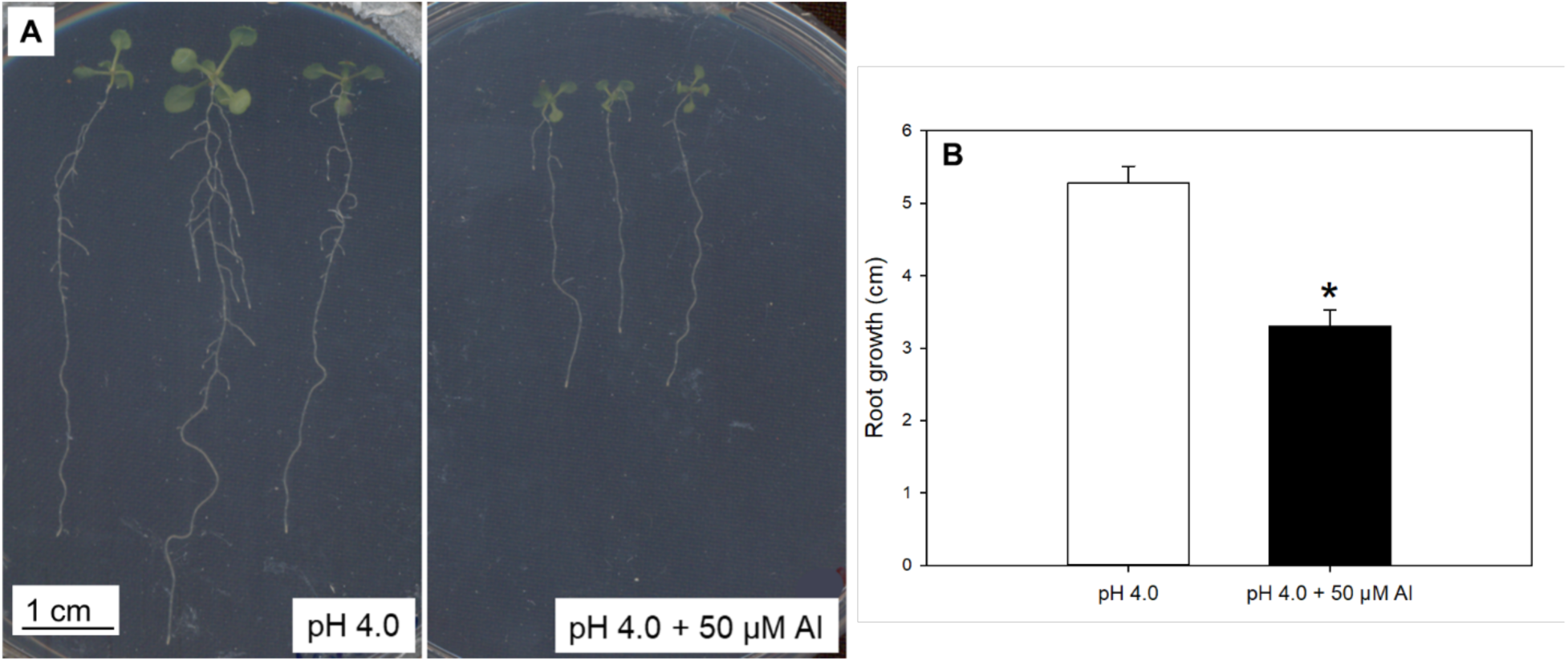
Aluminium (Al) Inhibits Root elongation in *Arabidopsis thaliana*. (a) Representative images of *Arabidopsis thaliana* seedlings (Columbia background, Col-0) scanned using the HP Scanjet G2410 after 10 days growth on MS medium plates without and with Al (50 µM AlCl_3_). (b) Root growth data were analysed after 10 days of exposure to a growth medium at pH 4.0 either without Al (- Al, 0 μM) or with Al (+ Al, 50 μM). Results are expressed as the mean ± standard error (n = 10). Image analysis was performed using ImageJ software. An asterisk (*) denotes a statistically significant difference between treatments for each genotype (Student *t*-test, \**p*<0.05).

### The TCA cycle is globally reprogrammed at the molecular and metabolic levels in response to Al

To access the translatome of the whole root, the cortical region of the elongation and maturation zones and the quiescent centre and endodermis regions, lines expressing *p35S:GFP-FLAG-RPL18*, *pPEP:GFP-FLAG-RPL18*, and *pSCR:GFP-FLAG-RPL18*, respectively, were used. Our results demonstrate that, in response to Al, mitochondrial metabolism is strongly tuned at a level of gene expression regulation that is very little explored and in specific cells that are very little accessed, which provides greater resolution and immeasurable contribution to the understanding of the underlying mechanisms associated with Al tolerance. Thus, the genes associated with the TCA cycle *ACONITASE* (*ACO*), *CITRATE SYNTHASE* (*CSY4*) and *FUMARASE* (*FUM*) were upregulated in the presence of Al and at the whole root level, with *ACO* and *CSY4* standing out, showing approximately 500- and 600-fold expression, respectively, compared to their respective controls (Figure 2 A, B and C). These results reinforce the mechanism in which OA produced in mitochondria-related reactions, such as citrate, malate and oxaloacetate, bind to and neutralise Al cations (Nunes-Nesi et al., 2014; Kochian et al., 2015), allowing root growth. To confirm these results, metabolite profiling at the whole root level was carried out. Thus, we found increases in the levels of TCA cycle intermediates in response to Al, such as malate, citrate and succinate, regardless of the genotype evaluated (Figure 3), corroborating the results of gene expression analyses at the whole root level.

**Figure 2.**
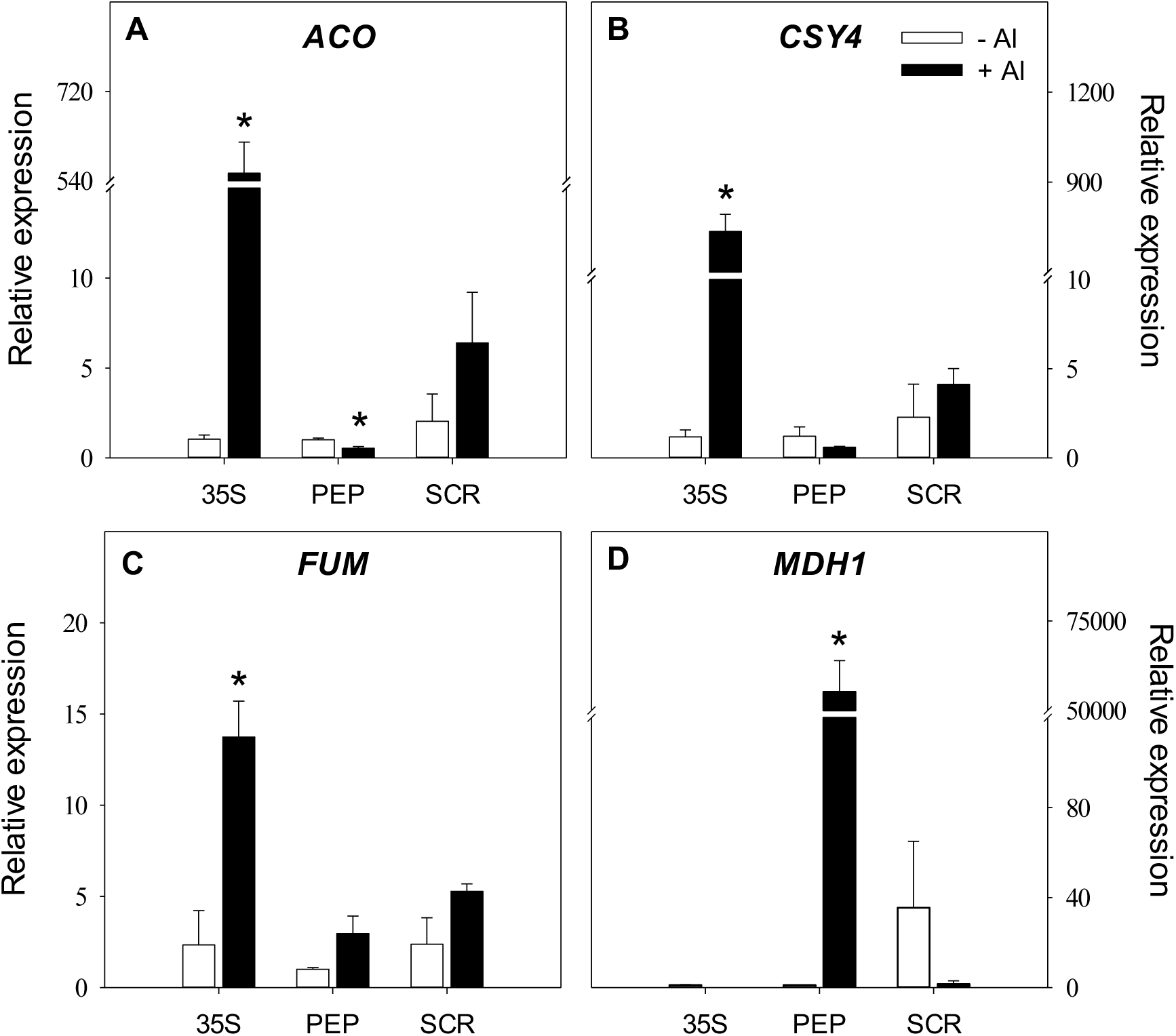
The TCA cycle is reprogrammed at translatome level and in specific cells in response to aluminium (Al). *Arabidopsis thaliana* seedlings (Columbia background, Col-0) expressing the *p:GFP-FLAG-RPL18* construct were used in this study, specifically: Cauliflower mosaic virus 35S (*p35S:GFP-FLAG-RPL18*), almost constitutive; plastid endopeptidase (*pPEP: GFP-FLAG-RPL18*), specific to the root cortex of the elongation and maturation zones; SCARECROW (*pSCR:GFP-FLAG-RPL18*), specific to the quiescent centre and root endodermis. After 10 days in growth medium without Al (- Al, 0 μM) or with Al (+ Al, 50 μM AlCl_3_), whole roots from the *p35S:GFP-FLAG-RPL18* line and whole seedlings from the *pPEP:GFP-FLAG-RPL18* and *pSCR:GFP-FLAG-RPL18* lines were collected at the start of the light period. Gene expression analysis was carried out according to the procedures described in Materials and Methods. Data were expressed as the mean between the values of 2^-(ΔΔCt)^ ± standard error (n = 3). An asterisk (*) denotes a statistically significant difference between treatments within each genotype (Student *t*-test, \**p*<0.05). Abbreviations: (a) *ACO*, *ACONITASE 1*; (b) *CSY*, *CITRATE SYNTHASE 4*; (c) *FUM*, *FUMARASE 1*; (d) *MDH*, *MALATE DEHYDROGENASE 1*.

**Figure 3.**
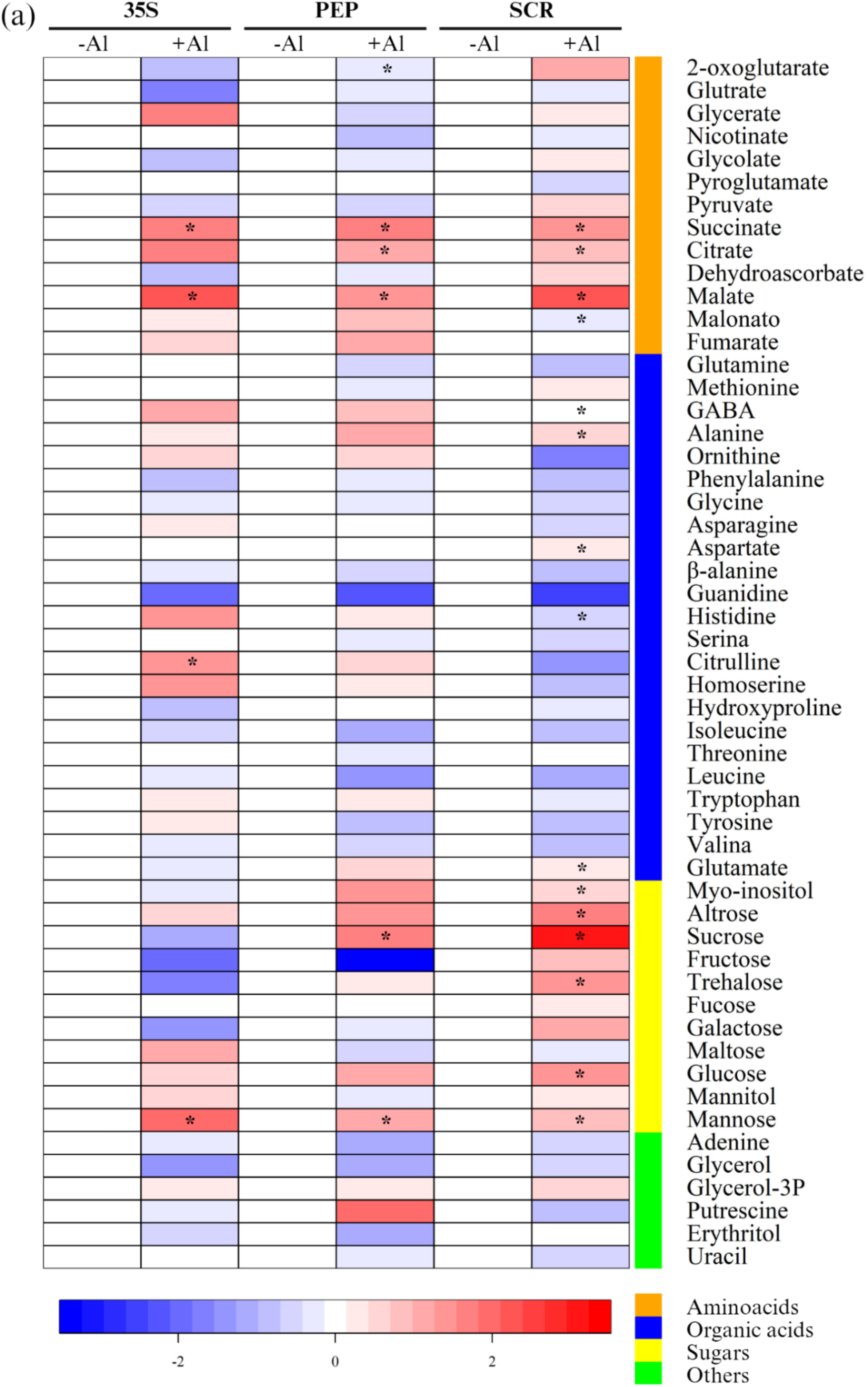
Metabolite profile is affected in response to aluminium (Al). *Arabidopsis thaliana* seedlings (Columbia background, Col-0) expressing the *p:GFP-FLAG-RPL18* construct were used in this study: Cauliflower mosaic virus 35S (*p35S:GFP-FLAG-RPL18*), almost constitutive; plastid endopeptidase (*pPEP: GFP-FLAG-RPL18*), specific to the root cortex of the elongation and maturation zones; SCARECROW (*pSCR:GFP-FLAG-RPL18*), specific to the quiescent centre and root endodermis. After 10 days in growth medium without Al (- Al, 0 μM) or with Al (+ Al, 50 μM AlCl_3_), whole roots of the *p35S:GFP-FLAG-RPL18, pPEP:GFP-FLAG-RPL18* and *pSCR:GFP-FLAG-RPL18* lines were collected at the start of the light period and promptly stored until metabolic profiling was carried out according to the procedures described in Materials and Methods. Metabolite profile data were expressed as the mean of four replicates, normalized in relation to the values found for each genotype grown in the absence of aluminium and log_2-_transformed for representation on the heat map. An asterisk (*) denotes a statistically significant difference between treatments within each genotype (Student *t*-test, \**p*<0.05).

### The *MDH* gene shows a differential distribution pattern between the meristematic and root differentiation regions in response to Al

Previous analyses of metabolic flux control have indicated that MALATE DEHYDROGENASE (MDH1) plays a major role in regulating the TCA cycle and that non-cyclic fluxes probably operate under stress conditions (Araújo et al., 2012). Notably, our work showed that the *MDH1* gene was upregulated in the cortical region of the elongation and maturation zones in response to Al, with expression approximately 53,000 times higher than in the control (Figure 2D). In addition, the reduced expression of *ACO* in the cortical region in response to Al (Figure 2A) suggests that citrate and especially malate are predominantly exuded to neutralise Al, reducing the carbon flow into the TCA cycle from citrate. Root OA exudation (citrate, malate and oxaloacetate) is associated with Al exclusion, where these exudates probably chelate Al in the soil (Kochian et al., 2015).

In acidic soils, emerging evidence suggests that the regulation of energy status and cell division involves unique interactions that modulate growth and detoxification responses (Balzergue et al., 2017; Mora-Macias et al., 2017; Ramesh et al., 2018; Hamasaki et al., 2019; Upadhyay et al., 2019; Duan et al., 2020; Siqueira et al., 2021). Under normal conditions, *Arabidopsis thaliana* roots treated exogenously with citrate, α-ketoglutarate and aconitate had a significantly reduced number of meristematic cells, while isocitrate and succinate had the opposite effect, significantly increasing the number of these cells (Zhang et al., 2023). Thus, TCA cycle metabolites lead to distinct root phenotypes in terms of stem cell division and growth. Furthermore, these phenotypes typically correlate with the localisation of these metabolites in the root. For example, succinate, located in the meristem, is a promoter of meristem growth, while aconitate, which is typically enriched in the differentiation zone, inhibits meristem growth and promotes cell elongation. Aconitate was the only metabolite that increased cell elongation, resulting in longer differentiated cells (Zhang et al., 2023). Although a trend towards increased expression of the *ACO*, *CSY4* and *FUM* genes was observed in the quiescent centre region in response to Al (Figure 2A, B and C), it is interesting to note that the *MDH1* gene was downregulated in this region in response to Al (Figure 2D). Thus, while endogenous malate homeostasis around the quiescent centre appears to be essential for root growth under Al exposure (Siqueira et al., 2022), our results point to a notable disturbance in this balance between the cortical and quiescent centre regions in response to Al exposure (Figure 2D).

### Cell cycle progression in meristematic cells, unlike that observed at the whole root level, is blocked in response to Al

Al-induced DNA damage activates the ATM and ATR kinases, which mediate the activation of the SOG1 transcription factor, inducing genes involved in DNA repair and cell cycle arrest. Together, ATM, ATR and SOG1 constitute an Al response pathway related to DNA damage caused by this element. Our results show an increase in ATR gene expression in the presence of Al and at whole root level, with few differences being observed in the cortical and quiescent centre regions in response to this element (Figure 4A). In contrast, a significant reduction in *ATM* gene expression was observed in the presence of Al and the quiescent centre region, while few differences were noted in the cortical region, as well as in the whole root, in response to Al (Figure 4B). These results, at least partially, must consider that *ATM*, *ATR* and *SOG1* exhibit an expression pattern that varies throughout the day, with a peak near dusk, which demonstrates the relevance of photoperiod to induce the DNA checkpoint response (Siqueira et al., 2021) and makes it possible to understand the reduced expression of *ATM* and *ATR*, especially in the quiescent centre region, in response to Al. Furthermore, in non-vegetable models, although ATR is essential in responses under conditions of DNA replication fork blockage, its activity can occur independently of DNA checkpoint or roles as a kinase, acting in mitochondria for example, where it prevents the release of cytochrome *c* into the cytosol and apoptosis (Hilton et al., 2015), which encourages us to hypothesise new roles for this protein in plant models, as suggested by the increase in *ATR* gene expression in the presence of Al and at whole root level found in this study. Meanwhile, the higher expression of the *ATM* gene in the absence of Al and the region of the quiescent centre may be associated with the interaction between ATM and the RCC1-UVR8-GEF-Like 3 (RUG3) protein to promote the functionality of mitochondrial complex I and the proliferation of root meristem cells, demonstrating the possible dual role of this protein in inducing the DNA damage response pathway and mitochondrial metabolic responses (Su et al., 2017).

**Figure 4.**
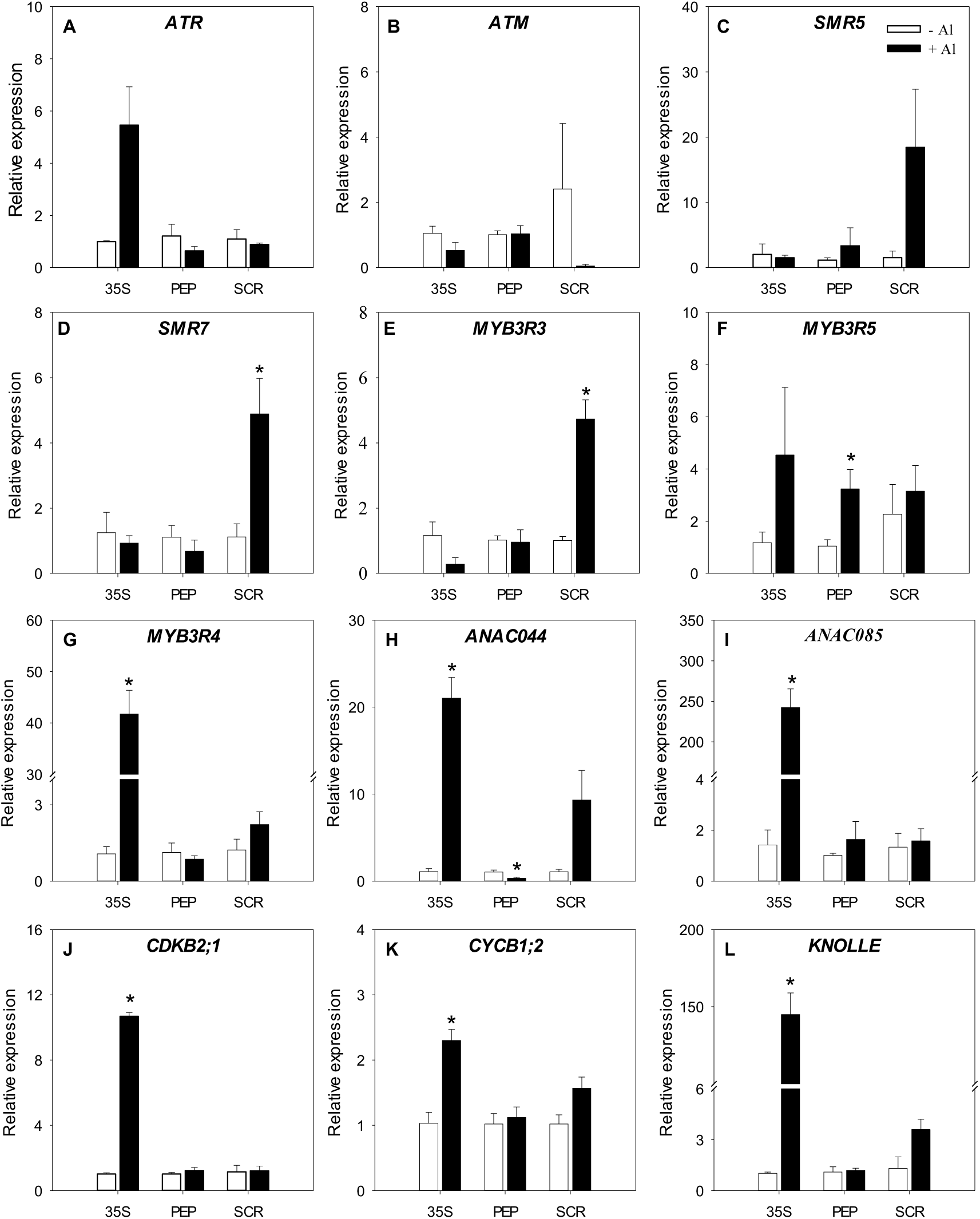
DNA damage response pathway is triggered at translatome level and in specific cells in response to aluminium (Al). *Arabidopsis thaliana* seedlings (Columbia background, Col-0) expressing the *p:GFP-FLAG-RPL18* construct were used in this study: Cauliflower mosaic virus 35S (*p35S:GFP-FLAG-RPL18*), almost constitutive; plastid endopeptidase (*pPEP: GFP-FLAG-RPL18*), specific to the root cortex of the elongation and maturation zones; SCARECROW (*pSCR:GFP-FLAG-RPL18*), specific to the quiescent centre and root endodermis. After 10 days in growth medium without Al (- Al, 0 μM) or with Al (+ Al, 50 μM AlCl_3_), whole roots from the *p35S:GFP-FLAG-RPL18* line and whole seedlings from the *pPEP:GFP-FLAG-RPL18* and *pSCR:GFP-FLAG-RPL18* lines were collected at the start of the light period. Gene expression analysis was carried out according to the procedures described in Materials and Methods. The data relating to gene expression analysis were expressed as the mean between the values of 2^-(ΔΔCt)^ ± standard error (n = 3). An asterisk (*) denotes a statistically significant difference between treatments within each genotype (Student *t*-test, \**p*<0.05). Abbreviations: (a) *ATR*, *ATAXIA TELANGIECTASIA MUTATED AND RAD3-RELATED*; (b) *ATM*, *ATAXIA TELANGIECTASIA MUTATED*; (c) *SMR5*, *SIAMESE-RELATED 5*; (d) *SMR7*, *SIAMESE-RELATED 7*; (j) *CDKB2;1*, *B2;1-TYPE CYCLIN-DEPENDENT KINASE*; (k) *CYCB1;2, B1;2-TYPE CYCLIN*.

Under conditions of DNA damage, CDK activity is inhibited, presumably by SMR5 and SMR7, transcriptionally activated by SOG1, resulting in the accumulation of Rep-MYB3Rs (Chen et al., 2017). Our results show higher expressions of the *SMR5* and *SMR7* genes in the presence of Al and the quiescent centre region, while smaller variations were observed in the cortical region and the whole root in response to the genotoxic agent (Figure 4C and D), indicating that the actions of the SMR5 and SMR7 proteins are predominantly confined to a specific group of cells. In fact, in the root apical meristem, the quiescent centre ensures DNA integrity by maintaining genome identity and preventing the spread of DNA damage to cells derived from the meristem.

In line with the higher expression of the *SMR5* and *SMR7* genes in the quiescent centre region in response to Al, we found an increase in the expression of the *MYB3R3* gene in this region as a function of Al (Figure 4E). On the other hand, greater expression of the *MYB3R5* gene in response to Al was demonstrated in the cortical region (Figure 4F). Taken together, these results suggest that although MYB3R3 and MYB3R5 share the same function, transcriptionally repressing genes related to the regulation of the transition between the G2 and M phases of the cell cycle and necessary for the onset of the M phase (Bourbousse et al., 2018), this function appears to be coordinated in a cell-specific manner, with MYB3R3 and MYB3R5 acting in a pronounced manner in the quiescent centre region and in the cortical region, respectively. In line with this coordinated cell-specific action of the MYB transcription factors, we found no changes in the expression of the *MYB3R4* gene in the quiescent centre and cortical regions in response to Al, but a notable increase in the expression of this gene in other cells (not identified in this study) in response to Al (Figure 4G). Chen et al. (2017), when examining MYB3R4 transcript and protein levels after treatment with roscovitine, a specific CDK inhibitor, found that they were elevated for some unknown reason. While our data provide new clues supporting an interaction between CDK and MYB3R4 proteins (as explained below), an alternative mechanism that involves CDK for MYB3R4 activation, as well as new findings about the functional roles of MYB3R4 under conditions of Al-induced DNA damage remain to be elucidated.

ANAC044 and ANAC085 are direct targets of SOG1, and the *anac044* and *anac085* knockout mutants are defective in inhibiting root growth under DNA stress, suggesting that ANAC044 and ANAC085 are responsible for the SOG1-dependent cell cycle pause (Takahashi et al., 2019). Our data reveal an increase in the expression of *ANAC044* and, above all, *ANAC085* at the whole root level in response to Al (Figure 4H and I). In addition, *ANAC044* expression was positively and negatively regulated in the quiescent centre and cortical regions, respectively, in the presence of Al (Figure 4H). Although our data do not support the involvement of ANAC044 and ANAC085 in the stabilisation of Rep-MYB3Rs proteins (Takahashi et al., 2019), they clearly demonstrate an involvement of ANAC044 and ANAC85 in the quiescent centre region and, more forcefully, in the other cells under Al exposure. In fact, considering that ANAC044 and ANAC085 are induced not only by DNA damage, but also by heat, cold and salt stress (Takahashi et al., 2019), this greater expression of ANAC044 and ANAC085 in other root cells suggests the involvement of these genes in other mechanisms that will culminate in plant tolerance to Al, in addition to those involved in the DNA damage-checkpoint-repair module.

Genotoxicity is implicated in DNA damage, affecting genome integrity and jeopardising DNA replication, which pauses growth by blocking cell cycle progression (Hu et al., 2016). Notably, this blockade usually occurs as a result of CDK/CYC inhibition at specific phases of the cell cycle, such as during the G1-S, intra-S and G2-M checkpoints (Hu et al., 2016). Our results showed little or no change in the expressions of the *CDKB2;1* and *CYCB1;2* genes in the quiescent centre and cortical regions in response to Al, in contrast to what was observed at the whole root level, where the expressions of these genes were upregulated in response to metal exposure (Figure 4J and K). CDK is known to phosphorylate and activate Act-MYBs, such as MYB3R4 (Araki et al., 2004; Haga et al., 2007). Thus, the lower expression of the *CDKB2;1*, *CYCB1;2* and *MYB3R4* genes at quiescent centre and cortical regions in response to Al leads us to theorize that the genotoxicity caused by exposure to this element at a concentration of 50 μM was responsible for pausing the cell cycle and was therefore sufficient to penalise the growth of the organ (Figure 1 and 4). Furthermore, the G2/M-specific *KNOLLE* gene, a target of repression by MYB3R3 and activation by MYB3R4, showed a pronounced increase in its expression at whole root level in response to Al (Figure 4L), suggesting a cell cycle with greater progression and low action of the MYB3R3 protein at whole root level. Al promoted greater MYB3R3 expression in the quiescent centre region, as demonstrated earlier in this study (Figure 4E).

## DISCUSSION

The elucidation of mechanisms regulating root growth has relied extensively on the visualization of transcriptional and translational reporters along root cell-types. Looking for a high-resolution model for reprogramming gene expression under sub-optimal conditions, two aspects must be considered. Firstly, several studies have demonstrated quantitative differences between the transcriptome (the pool of total mRNAs) and the translatome (ribosome-associated mRNAs) of plants, especially during stress conditions (; Piques et al., 2009; Poidevin et al., 2020; Li et al., 2021 Sainz et al., 2024). Secondly, studies of gene expression in specific cell types are greatly limited by the difficulties of isolating them without substantial contamination from other cells or neighbouring tissues (King and Gerber, 2016). The translatome of specific root cells is a promising tool for unravelling how mitochondrial metabolism, essentially the TCA cycle, and the DNA damage response (DDR) pathway, with special emphasis on cell cycle regulation, are remodelled in cells with distinct functions, particular patterns of gene expression, and differential responses to various types of stress. Thus, lines expressing *p35S*, *pPEP*, and *pSCR* were used to assess metabolic and DNA checkpoints at the translational level of gene expression in the whole root, the quiescent centre region, and the cortical region, respectively, in response to Al exposure. This contributes to elucidation of the internal cell-specific Al detoxification mechanism in plants.

Different genotypes can show contrasting metabolic responses to Al tolerance, with variations in OA levels being particularly remarkable (Pereira-Lima et al., 2023). Notably striking were the differential gradients observed in the normalized intensity of TCA cycle metabolites along the maize root development axis: succinate was most intense in the meristem, while aconitate, malate, and fumarate were typically enriched towards the root differentiation region (Zhang et al., 2023). In particular, the expression levels of the genes *ACO*, *SUCCINATE*-*COENZYME A LIGASE*, *SUCCINATE DEHYDROGENASE*, and *FUM* were correlated with the distribution of metabolites observed in maize roots (Zhang et al., 2023). Accordingly, for *A. thaliana*, spatially distinct distribution patterns of several classes of TCA cycle biosynthetic genes were found (Zhang et al., 2023). Our results demonstrate the reprogramming of the TCA cycle at the whole root level in response to Al exposure. The *ACO*, *CSY4*, and *FUM* genes were upregulated, while the *MDH1* gene was downregulated in response to this genotoxic agent (Figure 2D). Surprisingly, the *ACO* and *CSY4* genes showed an approximate 500-fold and 600-fold increase in expression, respectively, while the expression of the *MDH1* gene was reduced by approximately 580-fold at the whole root level in response to Al (Figure 2A-D). Therefore, the enhanced operation of the TCA cycle in the presence of Al can be attributed to the increased root demand for malate, which is involved in the internal detoxification of Al. When we zoomed in on the root, reaching the cortical region of the elongation and maturation zones, we impressively saw an approximately 53,000-fold increase in the expression of the *MDH1* gene in response to the presence of Al (Figure 2D). MDH exhibits a more linear correlation with respiration, triggering non-cyclic TCA cycle fluxes under stress (Araújo et al., 2012). As mentioned, Al detoxification by OAs can occur internally. The NIP1;2 transporter has been shown to facilitate the transport of Al from the root to the shoot, but only when it is in the form of an Al-malate complex (Wang et al., 2017), reinforcing the importance of intracellular malate content for Al tolerance. Additionally, an increase in intracellular malate levels was associated with a more tolerant phenotype, as observed for NADP-dependent malic enzyme (NADP-ME1) mutants (Badia et al., 2019). In this context, Al affects the plant cell cycle and inhibits root growth by altering the differentiation process of stem cells (Eekhout et al., 2017; Horvath et al., 2017), with the root distal transition zone (DTZ) considered the region most susceptible to Al toxicity (Kopittke et al., 2015). This susceptibility could be directly linked to the significant upregulation of *MDH1* in the root cortex and maturation zones of plants exposed to Al treatment. Functional connections between mitochondrial bioenergetics and DNA repair along the transition from the G2-M phases of the cell cycle have recently been postulated in plants (Siqueira et al., 2018). The inhibition of specific mitochondrial reactions resulted in the downregulation of cell cycle-related genes involved in the G2-M transition (Jardim-Messeder et al., 2015). Similarly, the antimycin A induces the inhibition of complex III of the mitochondrial electron transport chain (mETC) promotes the interaction between the kinase complex SNF1-RELATED PROTEIN KINASE 1 (SnRK1), an energy sensor that can modulate the production of OAs, with SOG1 (Hamasaki et al., 2019).

Overexpression of the *ACO1* gene in the meristem using the CYCB1;1 promoter significantly decreased primary root length due to the lower number of meristematic cells and the shortening of mature cells when compared to the control, while overexpression of this gene in the differentiation zone using the SMR1 promoter significantly increased root growth when compared to the control due to the higher number of meristematic cells (Zhang et al., 2023). In addition, changing the location of ACO overexpression from the meristem to the differentiation zone resulted in a different response to phosphate deprivation stress, although not to osmotic stress. Overall, pCYC::ACO lines are more resistant to phosphate deprivation stress conditions compared to pSMR1::ACO lines (Zhang et al., 2023). Thus, modifying the distribution pattern of a single aconitate and oxaloacetate biosynthesis gene, involved in responses to phosphate deprivation stress and aluminium toxicity, respectively, is sufficient to alter root development and stress response.

To determine whether exogenous applications of TCA cycle metabolites affected the ATP balance in roots, Zhang et al. (2023) analysed the localization pattern and intensity of a fluorescent ATP sensor in response to changes in TCA cycle inputs. Important changes were induced by treatments with oxaloacetate and α-ketoglutarate, which increased the intensity of the ATP sensor in the meristem. Isocitrate was the only metabolite tested that significantly increased the intensity of the ATP sensor in the meristem and the differentiation zone. Thus, although our study proposes a differential regulation of mitochondrial energy status in response to Al, changes in ATP levels may not be sufficient to fully elucidate the differential phenotypes caused by Al treatment and mediated by variations in the expression levels of TCA cycle metabolite biosynthesis genes.

When we zoomed in on the quiescent centre region to elucidate how the DDR pathway, particularly cell cycle regulation, is orchestrated as a function of Al exposure, the increase in the expression of the *SMR5*, *SMR7*, *MYB3R3*, and *ANAC044* genes (Figure 4C, D, E, and H) demonstrates an overactive DNA checkpoint regulation in this region. Thus, SMRs act by blocking the degradation of MYB3R3, while ANAC044 acts by conferring stability to this transcription factor, which is crucial for the genes required for the transition between the G2 and M phases of the cell cycle to have their expression suppressed. Genes with functions related to the cell cycle are preferentially regulated under the dual control of the E2F and MYB3R transcription factors (Gombos et al., 2023). Furthermore, MYB3Rs only act on mitotic genes, which explains the maintenance of cell quiescence after the loss of these transcription factors, unlike, for example, the breakdown of quiescence after the loss of RETINOBLASTOMA-RELATED (RBR) and the three canonical E2Fs (E2FA, E2FB, and E2FC), since these regulate the G1/S and G2/M transitions of the cell cycle (Gombos et al., 2023). Thus, considering that the quiescent centre ensures DNA integrity by conserving genomic identity and preventing the transmission of damaged DNA to the cells derived from the meristem, a DNA damage checkpoint mechanism must stand out, pausing the cell cycle in response to this damage and ensuring DNA repair and the maintenance of genome integrity. It seems reasonable to assume that to ensure a functional DDR pathway, suppression of genes related to the transition between the G2, and M phases of the cell cycle is likely essential, inhibiting cell division in the roots (Figure 5).

**Figure 5.**
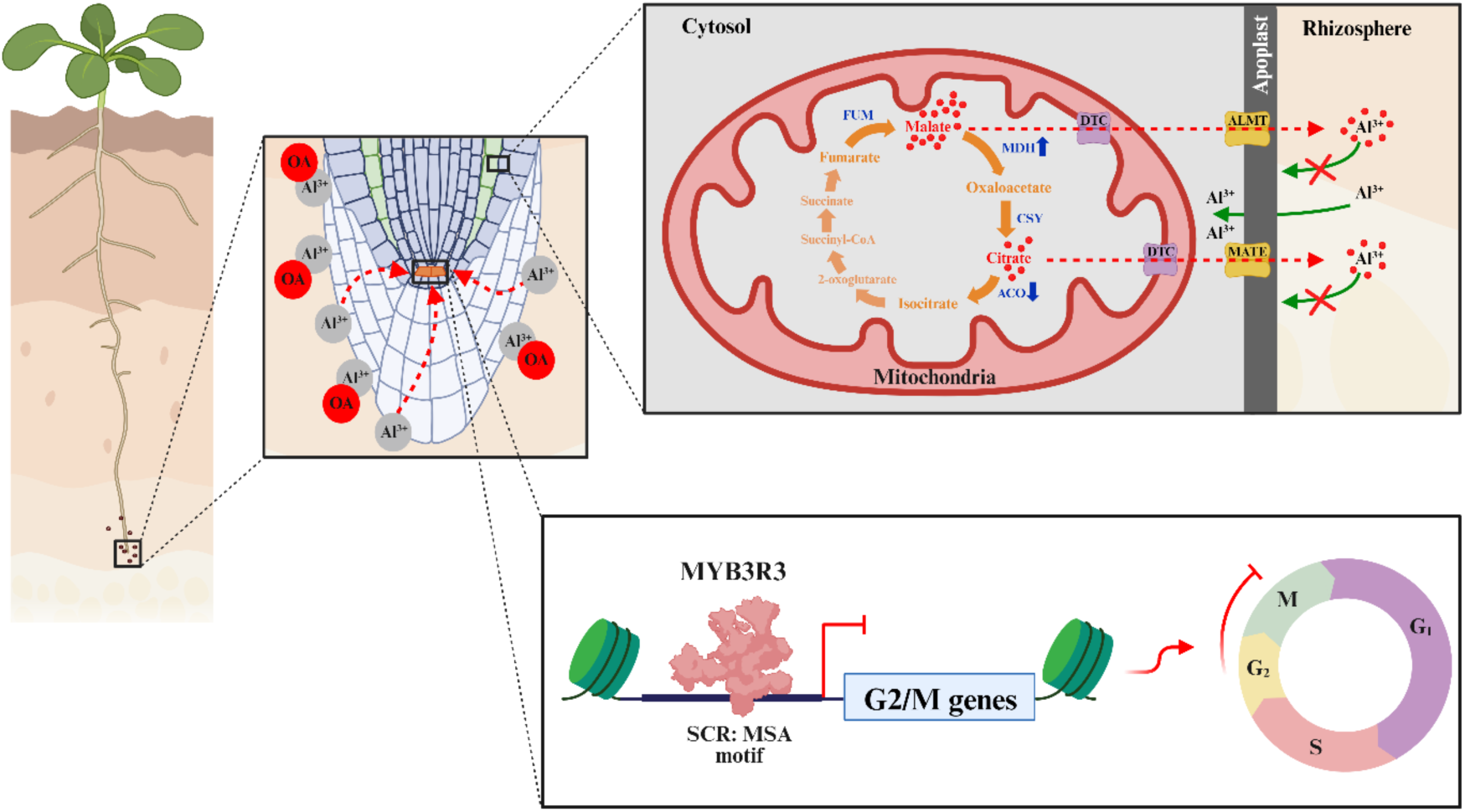
Coordination between the TCA cycle and the cell cycle in a cell-specific manner modulates aluminium toxicity responses in *Arabidopsis thaliana*. By analysing the translatome in specific cell populations, we observed a cell-specific coordination between DNA checkpoints and mitochondrial metabolism. Our results demonstrated that cell cycle progression, regulated by the MYB3R3 transcription factor, and mitochondrial bioenergetics associated with OA metabolism, particularly malate, occur primarily in the quiescent centre and the cortex of the elongation and maturation regions, respectively. Abbreviations: ACO, aconitase 1; Al^3+^, aluminum; ALMT, aluminum-activated malate transporter; CSY, citrate synthase 4; DTC, dicarboxylate/tricarboxylate carrier; FUM, fumarase 1; MATE, multidrug and toxic compound extrusion family; MDH, malate dehydrogenase 1; MSA, mitosis-specific activator; OA, organic acid; SCR, scarecrow.

By zooming in on the cortical region, we found less variation in the level of expression of the genes associated with cell cycle regulation in response to Al, except the increase in expression of the *MYB3R5* gene and the reduction in expression of the *ANAC044* gene (Figure 4F and H). Thus, while MYB3R3 is particularly concentrated in the region of the quiescent centre, MYB3R5 reveals a more distributed location among the populations of cell groups that make up the root. Focusing on the whole root, we found an increase in *ATR* gene expression in response to Al (Figure 3A). While ATM contributes to responses to DSBs, ATR responds to a wide range of events linked to DNA damage, including those that interfere with DNA replication (Yoshiyama et al., 2013), recruitment of RBR1 to sites of DNA damage to mediate the assembly of DNA-bound repair complexes (DRCs) (Bienderman et al., 2017; Horvath et al., 2017), and correction of single-strand breaks in DNA. In addition, ATR modulates quiescent centre differentiation under conditions of Al-dependent DNA damage (Zhang et al., 2018). Thus, the unchanged expression of the *ATR* gene in the quiescent centre region in response to Al suggests that the level of disturbance due to this element was not sufficient to induce cell differentiation in the region but allowed the maintenance of the quiescent centre to support root elongation under Al stress. Furthermore, we cannot rule out the relevance of photoperiod in the induction of DNA checkpoint responses to explain the low expression of *ATM* and *ATR* in the quiescent centre, since ATM, ATR, and SOG1 reach their peak expression around dusk (Siqueira et al., 2021). The increase in the expression of *ANAC044* and, substantially, *ANAC085* (Figure 4H and I) contrasted with the insignificant accumulation of Rep-MYB3Rs (Figure 4E and F) due to exposure to Al, suggests additional roles for these transcription factors beyond those associated with cell cycle checkpoints. Indeed, ANAC044 and ANAC085 are induced not only by DNA damage but also by heat, cold, and salinity (Takahashi et al., 2019). Further studies must provide new clues about the roles of these transcription factors in the presence of Al. Interestingly, the increase in the expression of the *CDKB2;1/CYCB1;2-MYB3R4* module and *KNOLLE* (Figure 4G, J, K and L), a target of repression by MYB3R3 and activation by MYB3R4, shows that DNA checkpoint regulation, especially that underlying cell cycle progression, is confined to specific groups of root cells, essentially the quiescent centre, providing support for cell cycle progression at the global level not being interrupted. Proteins relevant to cell cycle progression, such as CYCB1/CDK1, are localized in the mitochondrial matrix, promoting ATP synthesis in yeast (Wang et al., 2014). This most likely improves cellular bioenergetic efficiency for the transition between the G2 and M phases of the cell cycle, improving DNA repair and the survival of yeast cells under genotoxic conditions (Wang et al., 2014; Qin et al., 2015). Therefore, we cannot rule out the possibility that CDKB2;1 and CYCB1;2, in addition to stimulating an enhanced cell cycle progression, are contributing to improved mitochondrial bioenergetics at the whole root level in response to Al stress in *Arabidopsis thaliana*.

## CONCLUSIONS

Both cell cycle and mitochondrial bioenergetics play intrinsic roles in improving DNA repair and the survival of organisms under genotoxic conditions. Our results revealed that metabolic and genetic checkpoints seem to share similar regulatory characteristics under Al stress, allowing root elongation even under adverse conditions. Reports on heterotrophic yeast and animal cells indicate the presence of a synergistic network between enzymes related to malate/citrate metabolism and DNA repair proteins. By studying the translatome in populations of specific cell groups, we highlight a cell-specific coordination between DNA checkpoints and mitochondrial metabolism, as the regulation of cell cycle progression and mitochondrial bioenergetics related to OA metabolism, especially malate, occurred essentially in the cells of the quiescent centre and the cortex of the elongation and maturation regions, respectively. We further posit that the regulatory mechanisms underlying both cell cycle and metabolic checkpoints were able to preserve DNA integrity and promote internal Al detoxification, converging to allow root cell elongation. Thus, by providing greater resolution regarding the compartmentalization of the physiological processes associated with Al tolerance in plants, fundamentally the cell cycle and mitochondrial metabolism checkpoints, we identify more promising targets to be manipulated in the search for greater crop tolerance to Al, especially in acidic soils. In summary, we provide a valuable framework for improving plant resilience to Al stress, with potential applications in crop breeding and genetic engineering aimed at enhancing tolerance to challenging soil conditions.

## Acknowledgements

This work was supported by funding from the National Council for Scientific and Technological Development (CNPq-Brazil) [Grant 407276/2021–1 and 406455/2022– 8], and the Foundation for Research Assistance of the Minas Gerais State (FAPEMIG-Brazil Grant RED-00060-23), and the Agency for the Support and Evaluation of Graduate Education (CAPES) [Finance Code 001]. Scholarships granted by CAPES-Brazil to W.A.S. and T.W., CNPq to A.O.M and M.F.S. and research fellowships granted by CNPq to D.M.R., A.N.-N., and W.L.A. are gratefully acknowledged.

## Disclosers

The authors declare that there is no conflict of interest

## Authorś contribution

W.A.S., A.O.M., and W.L.A. designed the research. W.A.S., and A.O.M. performed most of the research; A.O.M and M.F.S. supervised the project; T.W., J.A.S., and D.B.M. contributed new reagents/analytic tools; D.M.R., A.R.F., and A.N-N. analyzed the data, discussed the results, and complemented the writing; W.A.S. and W.L.A. analyzed the data and wrote the article with input from all the others.

## Supplemental Material

**Supplementary Figure 1:**
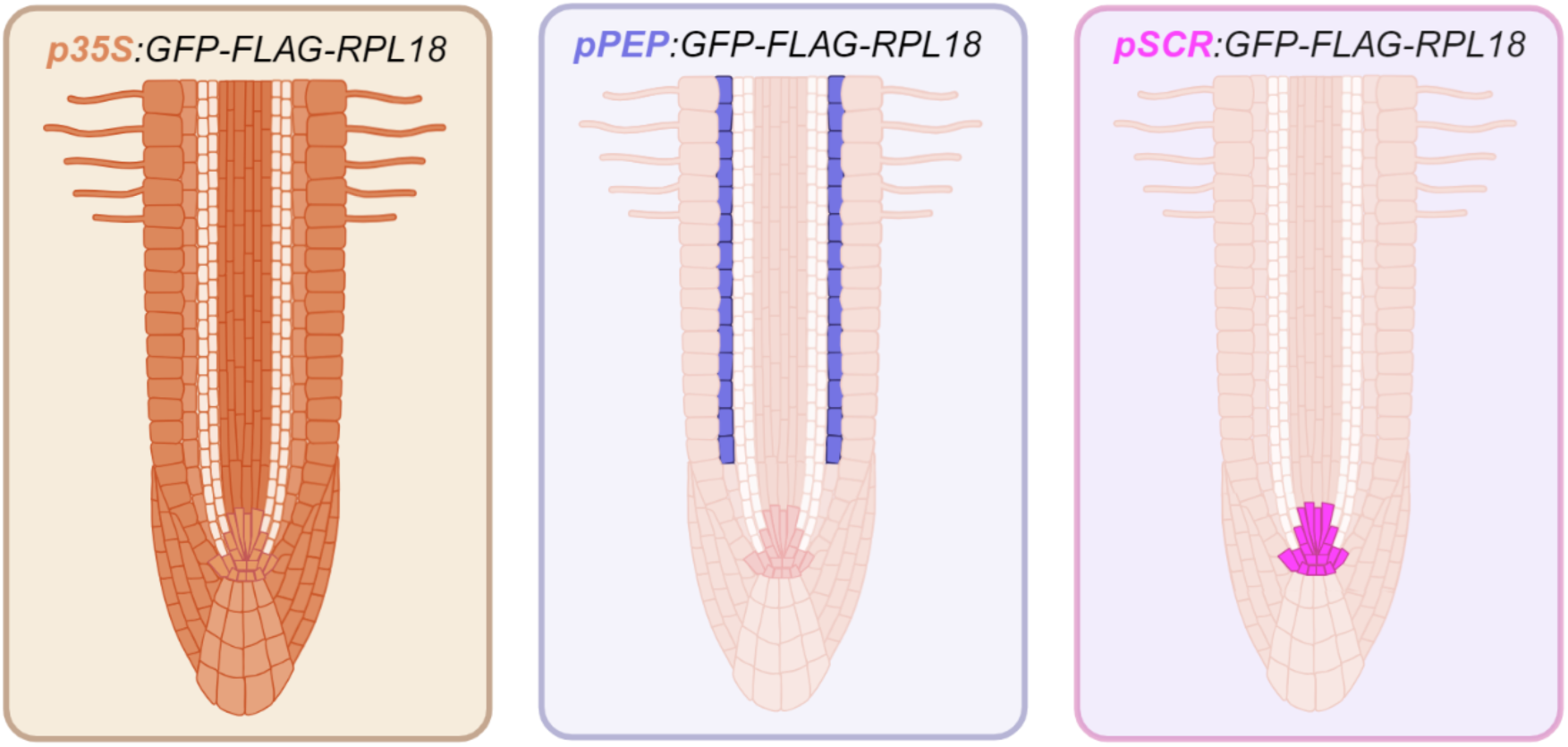
Schematic model of *Arabidopsis thaliana* translatome roots. Roots expressing the *p:GFP-FLAG-RPL18* construct (*p*: promoter; *GFP*: green fluorescent protein; *FLAG-RPL18*: FLAG-epitope tagged ribosomal protein L18). Individual promoters were used to direct the expression of the transgene, namely: Cauliflower mosaic virus 35S (*p35S:GFP-FLAG-RPL18*), almost constitutive; plastid endopeptidase (*pPEP:GFP-FLAG-RPL18*), specific to the root cortex of the elongation and maturation zones; SCARECROW (*pSCR:GFP-FLAG-RPL18*), specific to the quiescent centre and root endodermis.

**Supplementary Figure 2:**
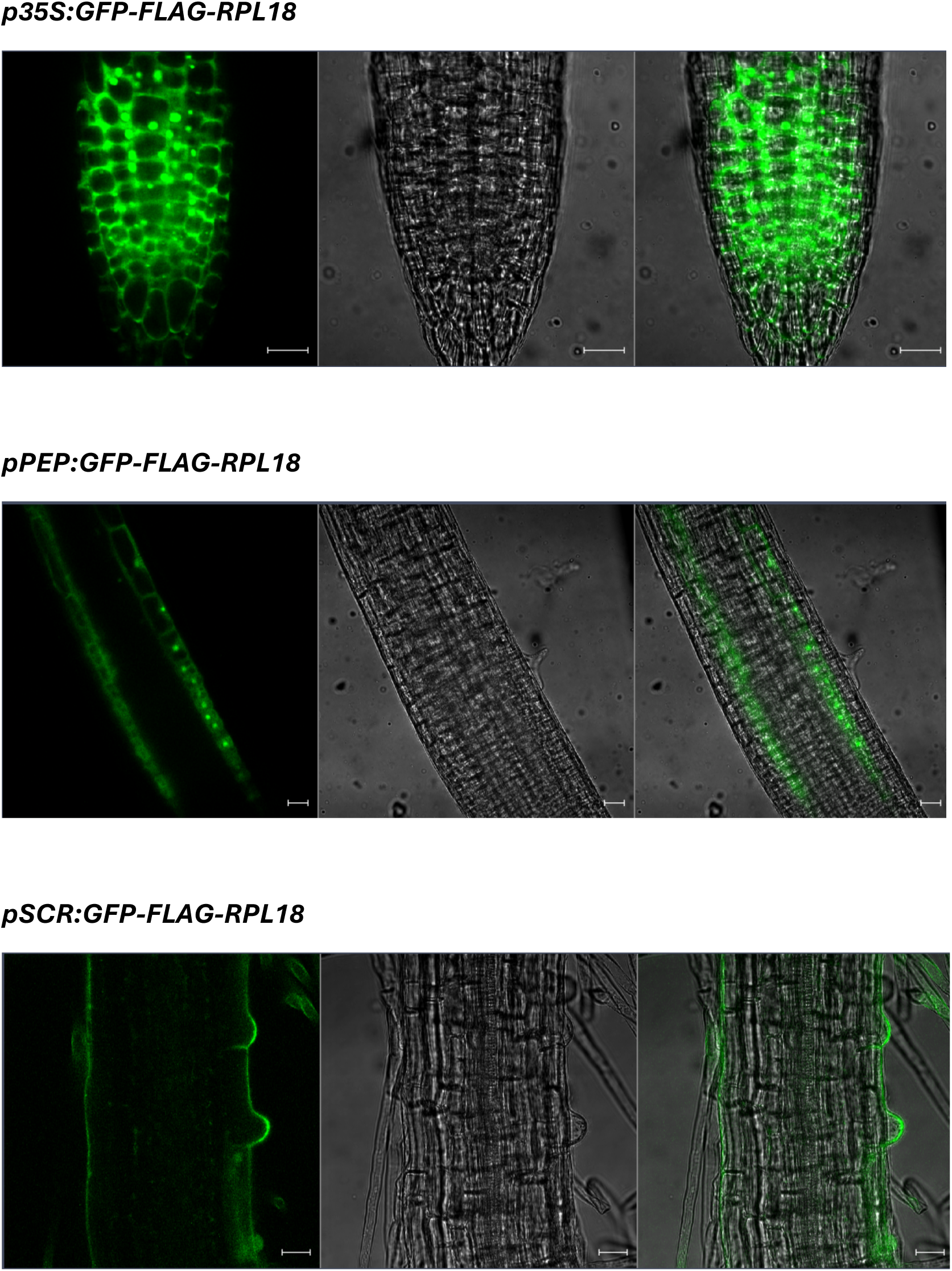
Confirmation of p:GFP-FLAG-RPL18 transgene expression. *Arabidopsis thaliana* seedlings expressing the p:GFP-FLAG-RPL18 construct: Cauliflower mosaic virus 35S (p35S:GFP-FLAG-RPL18), almost constitutive; plastid endopeptidase (pPEP: GFP-FLAG-RPL18), specific to the root cortex of the elongation and maturation zones; SCARECROW (pSCR:GFP-FLAG-RPL18), specific to the quiescent centre and root endodermis. Confirmation of the constructs was carried out using a Zeiss LSM510 META Laser Scanning Confocal Microscope using root apices excised from 7-day-old seedlings. The wavelengths used for excitation and emission of the GFP protein were 488 and 510 nm, respectively. The images were registered using ZEISS ZEN 3.1 software (scale bars = 20 μm).

**Supplementary Table 1:**
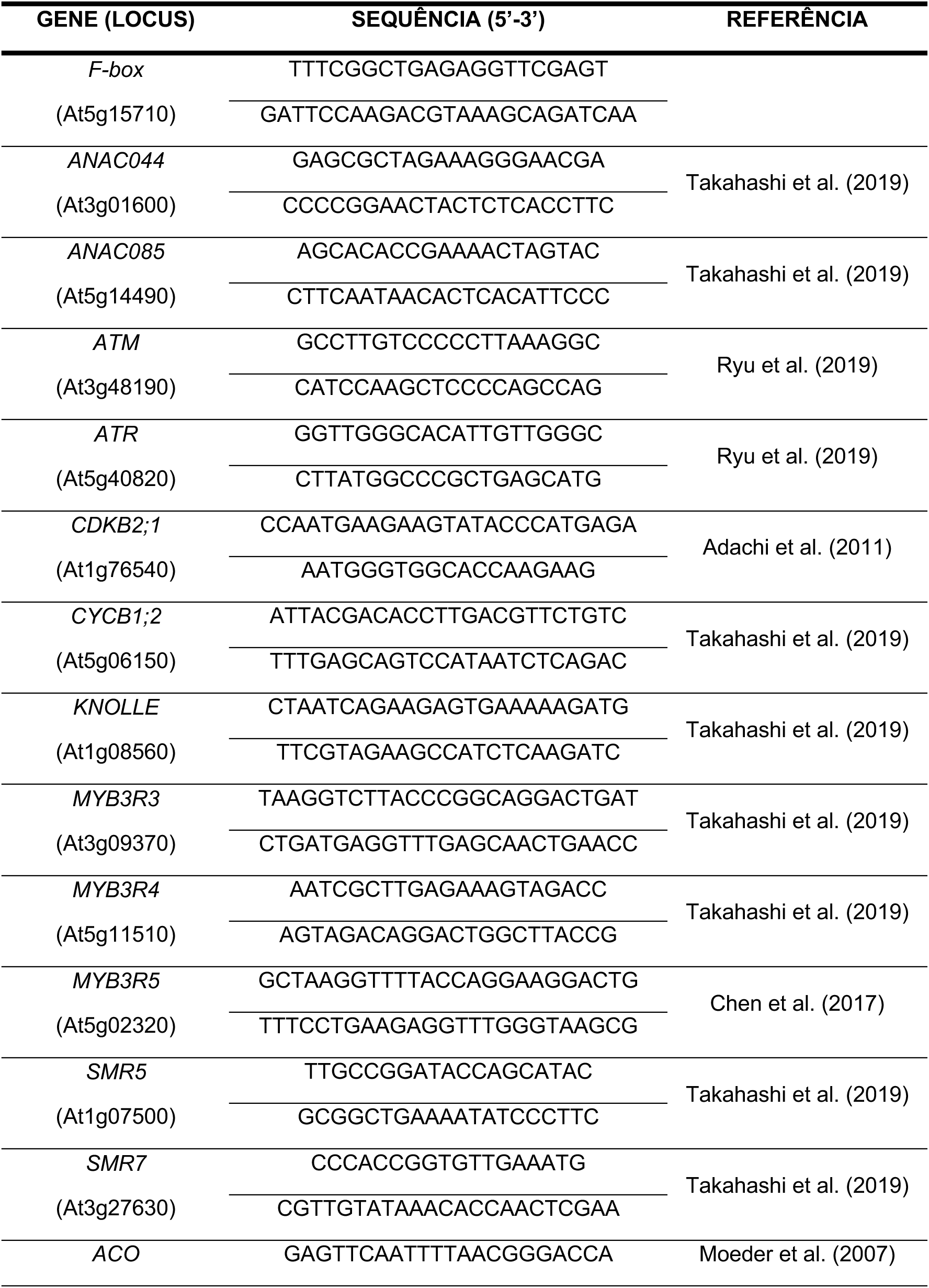
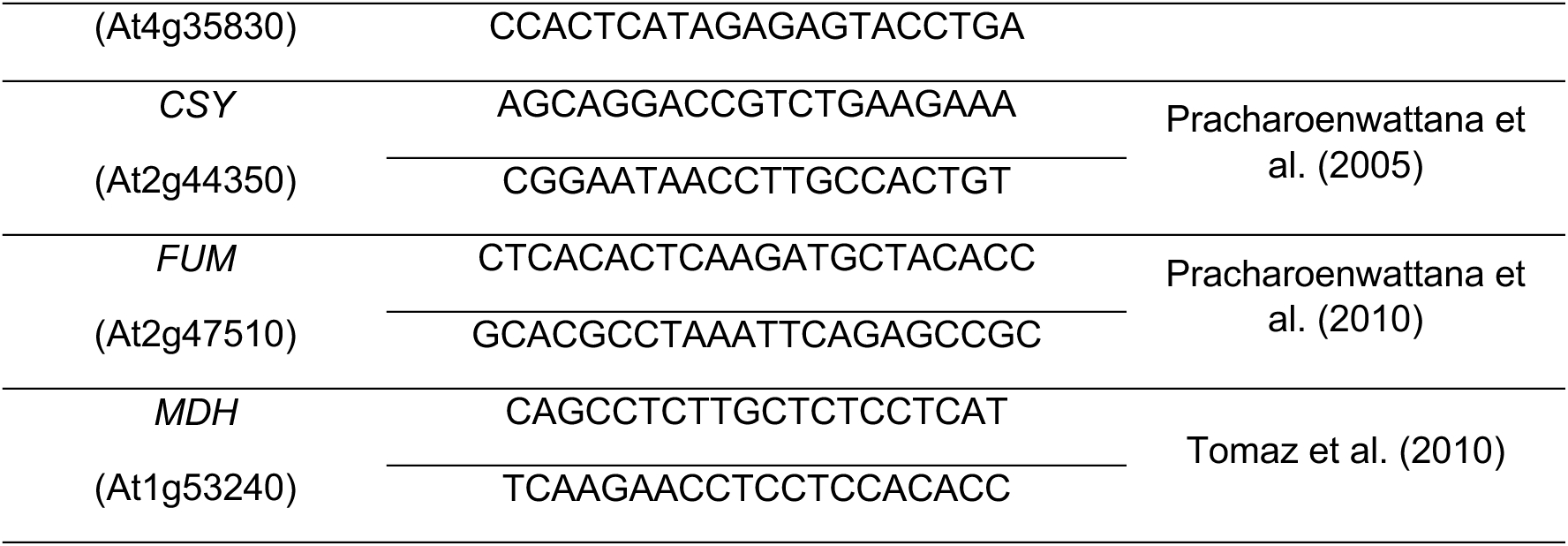
Primers used for qRT-PCR. Abreviations: ATM, ATAXIA TELANGIECTASIA MUTATED; ATR, ATAXIA TELANGIECTASIA MUTATED AND RAD3-RELATED; CDKB2;1, B2;1-TYPE CYCLIN-DEPENDENT KINASE; CYCB1;2, B1;2-TYPE CYCLIN; SMR5, SIAMESE-RELATED 5; SMR7, SIAMESE-RELATED 7; ACO, ACONITASE 1; CSY, CITRATE SYNTHASE 4; FUM, FUMARASE 1; MDH, MALATE DEHYDROGENASE 1.

## Notes

### Competing Interest Statement

The authors have declared no competing interest.

## REFERENCES

1. Adachi S, Minamisawa K, Okushima Y, Inagaki S, Yoshiyama K, Kondou Y, Kaminuma E, Kawashima M, Toyoda T, Matsui M, Kurihara D, Matsunaga S, Umeda M. 2011. Programmed induction of endoreduplication by DNA double-strand breaks in *Arabidopsis*. Proceedings of the National Academy of Sciences, USA 108, 10004– 10009.

2. Araki S, Ito M, Soyano T, Nishihama R, Machida Y. 2004. Mitotic cyclins stimulate the activity of c-Myb-like factors for transactivation of G2/M phase-specific genes in tobacco. The Journal of Biological Chemistry 279, 32979–32988.

3. Araújo WL, Nunes-Nesi A, Nikoloski Z, Sweetlove LJ, Fernie AR. 2012. Metabolic control and regulation of the tricarboxylic acid cycle in photosynthetic and heterotrophic plant tissues. Plant Cell and Environment 35, 1–21.

4. Badia MB, Maurino VG, Pavlovic T, Arias CL, Pagani MA, Andreo CS, Saigo M, Drincovich MF, Wheeler MCG. 2019. Loss of function of Arabidopsis NADP-malic enzyme 1 results in enhanced tolerance to aluminum stress. The Plant Journal 101, 653– 665.

5. Balzergue C, Dartevelle T, Godon C, Laugier E, Meisrimler C, Teulon J-M, Creff A, Bissler M, Brouchoud C, Hagège A, Müller J, Chiarenza S, Javot H, Becuwe-Linka N, David P, Péret B, Delannoy E, Thibaud M-C, Armengaud J, Abel S, Pellequer J-L, Nussaume L, Desnos T. 2017. Low phosphate activates STOP1-ALMT1 to rapidly inhibit root cell elongation. Nature Communications 8, 15300.

6. Bourbousse C, Vegesna N, Law JA. 2018. SOG1 activator and MYB3R repressors regulate a complex DNA damage network in *Arabidopsis*. Proceedings of the National Academy of Sciences, USA 115, 12453–12462.

7. Branco-Price C, Kaiser KA, Jang CJH, Larive CK, Bailey-Serres J. 2008. Selective mRNA translation coordinates energetic and metabolic adjustments to cellular oxygen deprivation and reoxygenation in *Arabidopsis thaliana*. The Plant Journal 56, 743–755.

8. Branco-Price C, Kawaguchi R, Ferreira RB, Bailey-Serres J. 2005. Genome-wide analysis of transcript abundance and translation in Arabidopsis seedlings subjected to oxygen deprivation. Annals of Botany 96, 647–660.

9. Chen P, Takatsuka H, Takahashi N, Kurata R, Fukao Y, Kobayashi K, Ito M, Umeda M. 2017. *Arabidopsis* R1R2R3-Myb proteins are essential for inhibiting cell division in response to DNA damage. Nature Communications 8, 635.

10. de Jong M, van Breukelen B, Wittink FR, Menke FLH, Weisbeek PJ, den Ackerveken GV. 2006. Membrane-associated transcripts in *Arabidopsis*; their isolation and characterization by DNA microarray analysis and bioinformatics. The Plant Journal 46, 708–721.

11. Duan S, Hu L, Dong B, Jin HL, Wang HB. 2020. Signaling from plastid genome stability modulates endoreplication and cell cycle during plant development. Cell Reports 32, 108019.

12. Eekhout T, Larsen P, De Veylder L. 2017. Modification of DNA checkpoints to confer aluminum tolerance. Trends in Plant Science 22, 102–105.

13. Fernie AR, Aharoni A, Willmitzer L, Stitt M, Tohge T, Kopka J, Carroll AJ, Saito K, Fraser PD, DeLuca V. 2011. Recommendations for reporting metabolite data. The Plant Cell 23, 2477–2482.

14. Fulcher N, Sablowski R. 2009. Hypersensitivity to DNA damage in plant stem cell niches. Proceedings of the National Academy of Sciences, USA 106, 20984–20988.

15. Furukawa T, Curtis MJ, Tominey CM, Duong YH, Wilcox BW, Aggoune D, Hays JB, Britt AB. 2010. A shared DNA-damage-response pathway for induction of stem-cell death by UVB and by gamma irradiation. DNA Repair 9, 940–948.

16. Gombos M, Raynaud C, Nomoto Y, Molnár E, Brik-Chaouche R, Takatsuka H, Zaki A, Bernula D, Latrasse D, Mineta K, Nagy F, He X, Iwakawa H, Őszi E, An J, Suzuki T, Papdi C, Bergis C, Benhamed M, Bögre L, Ito M, Magyar Z. 2023. The canonical E2Fs together with RETINOBLASTOMA-RELATED are required to establish quiescence during plant development. Communications Biology 6, 903.

17. Haga N, Kato K, Murase M, Araki S, Kubo M, Demura T, Suzuki K, Muller I, Voss U, Jurgens G, Ito M. 2007. R1R2R3-Myb proteins positively regulate cytokinesis through activation of *KNOLLE* transcription in *Arabidopsis thaliana*. Development 134, 1101– 1110.

18. Hamasaki H, Kurihara Y, Kuromori T, Kusano H, Nagata N, Yamamoto YY, Shimada H, Matsui M. 2019. SnRK1 kinase and the NAC transcription factor SOG1 are components of a novel signaling pathway mediating the low energy response triggered by ATP depletion. Frontiers in Plant Science 10, 503.

19. Hilton BA, Li Z, Musich PR, Wang H, Cartwright BM, Serrano M, Zhou XZ, Lu KP, Zou Y. 2015. ATR plays a direct antiapoptotic role at mitochondria, which is regulated by prolyl isomerase Pin1. Molecular Cell 60, 35–46.

20. Horvath BM, Kourova H, Nagy S, Nemeth E, Magyar Z, Papdi C, Ahmad Z, Sanchez-Perez GF, Perilli S, Blilou I, Pettkó-Szandtner A, Darula Z, Meszaros T, Binarova P, Bogre L, Scheres B. 2017. *Arabidopsis* RETINOBLASTOMA RELATED directly regulates DNA damage responses through functions beyond cell cycle control. The EMBO Journal 36, 1261–1278.

21. Hu Z, Cools T, De Veylder L. 2016. Mechanisms used by plants to cope with DNA damage. Annual Review of Plant Biology 67, 439–462.

22. Jardim-Messeder D, Caverzan A, Rauber R, de Souza FE, Margis-Pinheiro M, Galina A. 2015. Succinate dehydrogenase (mitochondrial complex II) is a source of reactive oxygen species in plants and regulates development and stress responses. New Phytologist 208, 776–789.

23. Kawaguchi R, Bailey-Serres J. 2005. mRNA sequence features that contribute to translational regulation in *Arabidopsis*. Nucleic Acids Research 33, 955–965.

24. Kawaguchi R, Girke T, Bray EA, Bailey-Serres J. 2004. Differential mRNA translation contributes to gene regulation under non-stress and dehydration stress conditions in *Arabidopsis thaliana*. The Plant Journal 38, 823–839.

25. King HA, Gerber AP. 2016. Translatome profiling: methods for genome-scale analysis of mRNA translation. Briefings in Functional Genomics 15, 22–31.

26. Kochian LV, Hoekenga AO, Piñeros MA. 2004. How do crop plants tolerate acid soils? Mechanisms of aluminum tolerance and phosphorus efficiency. Annual Review of Plant Biology 55, 459–493.

27. Kochian LV, Piñeros MA, Liu J, Magalhaes JV. 2015. Plant adaptation to acid soils: The molecular basis for crop aluminum resistance. Annual Review of Plant Biology 66, 571–598.

28. Kopittke PM, Moore KL, Lombi E, Gianoncelli A, Ferguson BJ, Blamey FPC, Menzies NW, Nicholson TM, McKenna BA, Wang P, Gresshoff PM, Kourousias G, Webb RI, Green K, Tollenaere A. 2015. Identification of the primary lesion of toxic aluminum in plant roots. Plant Physiology 167, 1402–1411.

29. Kopka J, Schauer N, Krueger S, Birkemeyer C, Usadel B, Bergmüller E, Dörmann P, Weckwerth W, Gibon Y, Stitt M, Willmitzer L, Fernie AR, Steinhauser D. 2005.: the golm metabolome database. Bioinformatics 21, 1635–1638.

30. Li Y, Li Q, Beuchat G, Zeng H, Zhang C, Chen LQ. 2021. Combined analyses of translatome and transcriptome in Arabidopsis reveal new players responding to magnesium deficiency. Journal of integrative plant biology, 63, 2075–2092.

31. Lisec L, Schauer N, Kopka J, Willmitzer L, Fernie AR. 2006. Gas chromatography mass spectrometry-based metabolite profiling in plants. Nature Protocols 1, 387–396.

32. Luedemann A, Strassburg K, Erban A, Kopka J. 2008. TagFinder for the quantitative analysis of gas chromatography-mass spectrometry (GC-MS)-based metabolite profiling experiments. Bioinformatics 24, 732–737.

33. Murashige T, Skoog F. 1962. A revised medium for rapid growth and bio assays with tobacco tissue cultures. Physiologia Plantarum 15, 473–497.

34. Mustroph A, Zanetti ME, Girke T, Bailey-Serres J. 2013. Isolation and analysis of mRNAs from specific cell types of plants by ribosome immunopurification. Methods in Molecular Biology 959, 277–302.

35. Mustroph A, Zanetti ME, Jang CJH, Holtan HE, Repetti PP, Galbraith DW, Girke T, Bailey-Serres J. 2009. Profiling translatomes of discrete cell populations resolves altered cellular priorities during hypoxia in *Arabidopsis*. Proceedings of the National Academy of Sciences, USA 106, 18843–18848.

36. Nicolaï M, Roncato MA, Canoy AS, Rouquié D, Sarda X, Freyssinet G, Robaglia C. 2006. Large-scale analysis of mRNA translation states during sucrose starvation in *Arabidopsis* cells identifies cell proliferation and chromatin structure as targets of translational control. Plant Physiology 141, 663–673.

37. Nunes-Nesi A, Brito DS, Inostroza-Blancheteau C, Fernie AR, Araújo WL. 2014. The complex role of mitochondrial metabolism in plant aluminum resistance. Trends in Plant Science 19, 399–407.

38. Ofoe R, Thomas RH, Asiedu SK, Wang-Pruski G, Fofana B, Abbey L. 2022. Aluminum in plant: Benefits, toxicity and tolerance mechanisms. Frontiers in Plant Science 13, 1085998.

39. Piques M, Schulze WX, Höhne M, Usadel B, Gibon Y, Rohwer J, Stitt M. 2009. Ribosome and transcript copy numbers, polysome occupancy and enzyme dynamics in *Arabidopsis*. Molecular Systems Biology 5, 314.

40. Poidevin L, Forment J, Unal D, Ferrando A. 2021. Transcriptome and translatome changes in germinated pollen under heat stress uncover roles of transporter genes involved in pollen tube growth. Plant, Cell & Environment, 44, 2167–2184.

41. Qin L, Fan M, Candas D, Jiang G, Papadopoulos S, Tian L, Woloschak G, Grdina DJ, Li JJ. 2015. CDK1 enhances mitochondrial bioenergetics for radiation-induced DNA repair. Cell Reports 13, 2056–2063.

42. Ramesh S, Kamran M, Sullivan W, Chirkova L, Okamoto M, Degryse F, McLaughlin M, Gilliham M, Tyerman SD. 2018. Split personality of aluminum activated malate transporter family proteins: facilitation of both GABA and malate transport. The Plant Cell 30, 1147–1164.

43. Schauer N, Steinhauser D, Strelkov S, Schomburg D, Allison G, Moritz T, Lundgren K, Roessner-Tunali U, Forbes MG, Willmitzer L, Fernie AR, Kopka J. 2005. GC-MS libraries for the rapid identification of metabolites in complex biological samples. FEBS Letters 579, 1332–1337.

44. Siqueira JA, Hardoim P, Ferreira PCG, Nunes-Nesi A, Hemerly AS. 2018. Unraveling interfaces between energy metabolism and cell cycle in plants. Trends in Plant Science 23, 731–747.

45. Siqueira JA, Silva MF, Wakin T, Nunes-Nesi A, Araújo WL. 2022. Metabolic and DNA checkpoints for the enhancement of Al tolerance. Journal of Hazardous Materials 430, 128366.

46. Siqueira JA, Wakin T, Batista-Silva W, Silva JCF, Vicente MH, Silva JC, Clarindo WR, Zsögön A, Peres LEP, De Veylder L, Fernie AR, Nunes-Nesi A, Araújo WL. 2021. A long and stressful day: Photoperiod shapes aluminium tolerance in plants. Journal of Hazardous Materials 432, 128704.

47. Sjogren CA, Bolaris SC, Larsen PB. 2015. Aluminum-dependent terminal differentiation of the Arabidopsis root tip is mediated through an ATR-, ALT2-, and SOG1-regulated transcriptional response. The Plant Cell 27, 2501–2515.

48. Su C, Zhao H, Zhao Y, Ji H, Wang Y, Zhi L, Li X. 2017. RUG3 and ATM synergistically regulate the alternative splicing of mitochondrial nad2 and the DNA damage response in Arabidopsis thaliana. Scientific Reports 7, 43897.

49. Takahashi N, Ogita N, Takahashi T, Taniguchi S, Tanaka M, Seki M, Umeda M. 2019. A regulatory module controlling stress-induced cell cycle arrest in *Arabidopsis*. eLife 8, e43944.

50. Upadhyay N, Kar D, Datta S. 2019. A multidrug and toxic compound extrusion (MATE) transporter modulates auxin levels in root to regulate root development and promotes aluminium tolerance. Plant Cell and Environment 43, 745–759.

51. Wang Y, Li R, Li D, Jia X, Zhou D, Li J. 2017. NIP1;2 is a plasma membrane-localized transporter mediating aluminum uptake, translocation, and tolerance in *Arabidopsis*. Proceedings of the National Academy of Sciences, USA 114, 5047–5052.

52. Wang Z, Fan M, Candas D, Zhang T-Q, Qin L, Eldridge A, Wachsmann-Hogiu S, Ahmed KM, Chromy BA, Nantajit D, Duru N, He F, Chen M, Finkel T, Weinstein LS, Li JJ. 2014. Cyclin B1/Cdk1 coordinates mitochondrial respiration for cell-cycle G2/M progression. Developmental Cell 29, 217–232.

53. Yi D, Kamei CLA, Cools T, Vanderauwera S, Takahashi N, Okushima Y, Eekhout T, Yoshiyama KO, Larkin J, Van den Daele H, Conklin P, Britt A, Umeda M, De Veylder L. 2014. The *Arabidopsis* SIAMESE-RELATED cyclin-dependent kinase inhibitors SMR5 and SMR7 regulate the DNA damage checkpoint in response to reactive oxygen species. The Plant Cell 26, 296–309.

54. Yoshiyama K, Conklin PA, Huefner ND, Britt AB. 2009. Suppressor of gamma response 1 (*SOG1*) encodes a putative transcription factor governing multiple responses to DNA damage. Proceedings of the National Academy of Sciences, USA 106, 12843– 12848.

55. Yoshiyama KO, Kobayashi J, Ogita N, Ueda M, Kimura S, Maki H, Umeda M. 2013. ATM-mediated phosphorylation of SOG1 is essential for the DNA damage response in *Arabidopsis*. EMBO Reports 14, 817–822.

56. Zhang T, Noll SE, Peng JT, Klair A, Tripka A, Stutzman N, Cheng C, Zare RN, Dickinson AJ. 2023. Chemical imaging reveals diverse functions of tricarboxylic acid metabolites in root growth and development. Nature Communications 14, 2567.

57. Zhang Y, Guo J, Chen M, Li L, Wang L, Huang C-F. 2018. The cell cycle checkpoint regulator ATR is required for internal aluminum toxicity-mediated root growth inhibition in *Arabidopsis*. Frontiers in Plant Science 9, 118

